# Sing to me, baby: Infants show neural tracking and rhythmic movements to live and dynamic maternal singing

**DOI:** 10.1101/2023.02.28.530310

**Authors:** Trinh Nguyen, Susanne Reisner, Anja Lueger, Samuel V. Wass, Stefanie Hoehl, Gabriela Markova

## Abstract

Infant-directed singing has unique acoustic characteristics that may allow even very young infants to respond to the rhythms carried through the caregiver’s voice. The goal of this study was to examine neural and movement responses to live and dynamic maternal singing in 7-month-old infants and their relation to linguistic development. In total, 60 mother-infant dyads were observed during two singing conditions (playsong and lullaby). In Study 1 (*n* = 30), we measured infant EEG and used an encoding approach utilizing ridge regressions to measure neural tracking. In Study 2 (*n* = 40), we coded infant rhythmic movements. In both studies, we assessed children’s vocabulary when they were 20 months old. In Study 1, we found above-threshold neural tracking of maternal singing, with superior tracking of lullabies than playsongs. We also found that the acoustic features of infant-directed singing modulated tracking. In Study 2, infants showed more rhythmic movement to playsongs than lullabies. Importantly, neural coordination (Study 1) and rhythmic movement (Study 2) to playsongs were positively related to infants’ expressive vocabulary at 20 months. These results highlight the importance of infants’ brain and movement coordination to their caregiver’s musical presentations, potentially as a function of musical variability.

---

Musicality, defined as the ability to perceive, process and produce music, is a human universal with deep ontogenetic roots (e.g., Mehr et al., 2021; Savage et al., 2021). Infants experience a variety of different types of music every day (Mendoza & Fausey, 2021; Warlaumont et al., 2022). Most prominently, caregivers sing to them to capture their attention, regulate their arousal, and share emotions with or entertain them (Trehub, 2019). On the other side, children have sophisticated perceptual capabilities when listening to music from early in life: they are sensitive to tempo- and pitch-related organization of musical sequences, such as changes in melodic patterns, harmony, grouping, beat, and meter (e.g., Ferland & Mendelson, 1989; Háden et al., 2015; Krumhansl & Jusczyk, 1990; Trainor & Heinmiller, 1998; Winkler et al., 2009). For example, frequency tagging studies show that infants’ brain activity increases relative to rhythmic beats in music, and that these increases are influenced by musical properties such as meter and by musical experience (Cirelli et al., 2016; Lenc et al., 2022). No research to date has, however, examined whether volume fluctuations in sung music directly map onto fluctuations in the brain activity in a listening child. Several recent studies have demonstrated similar correspondence between volume fluctuations in child-directed speech and children’s brain activity (Attaheri, Choisdealbha, Rocha, et al., 2022; Menn, Ward, et al., 2022), and so we were interested to investigate whether infants also show neural tracking of music.

In addition, we wanted to understand better the mechanisms that might subserve neural tracking of music. Here, we refer to neural tracking such that an exogenous signal and an endogenous activity are temporally aligned, also referred to as neural entrainment in the broad sense (Obleser & Kayser, 2019). In particular, we were interested to study the association between infants’ rhythmic movement to music (Sievers et al., 2013) and neural tracking. Fetuses at 35 weeks of gestational age already show more movement to music in comparison to speech (Kisilevsky et al., 2004), and by 2 months of age, they coordinate their gaze with the beat of infant-directed (ID) singing (Lense et al., 2022). During later infancy, they move their bodies more to music than to other auditory stimuli, and modulate the tempo of their movement contingent on the tempo of the music (Ilari, 2015; Zentner & Eerola, 2010). Because rhythmic movements are known to guide auditory perception (Phillips-Silver & Trainor, 2005, 2007), it has been suggested that this coordination, in particular to the rhythmic structure of the musical input, may modify brain activity and behavior in such a way that they eventually lock onto a common periodicity (Vuilleumier & Trost, 2015). However, no previous research has investigated this hypothesis. Previous discussions of neural tracking of language have concentrated on a range of different factors that drive language entrainment, such as understanding how temporal fluctuations in the critical band envelope of speech directly entrain brain activity (Doelling et al., 2014), and how cognitive factors such as the comprehensibility of speech mediate neural entrainment (Poeppel & Assaneo, 2020). In the present study we examined the association between neural tracking and rhythmic movement to provide a new perspective on the mechanisms that drive neural entrainment to music.

We also wanted to investigate the three-way relationship between neural tracking, rhythmic movements to music, and language development. Recent evidence suggests that neural tracking of nursery rhymes is predictive of infants’ later language outcomes (Attaheri, Choisdealbha, Rocha, et al., 2022; Menn, Ward, et al., 2022). Brandt and colleagues (2012) propose that the musical aspects of language (i.e., prosody, rhythm, timbral contrast) scaffold the later development of semantic and syntactic aspects of language. In fact, available evidence suggests that an enriched musical environment (i.e., music classes, music at home) during infancy and preschool can promote language development (Papadimitriou et al., 2021; Politimou et al., 2019; Putkinen et al., 2013; Williams et al., 2015; Zhao & Kuhl, 2016). Brain responses of newborns to sung as compared to spoken streams of syllables predict expressive vocabulary at 18 months (François et al., 2017), and ID singing, in particular, has been found to facilitate aspects of phonetic perception and word learning in infants in the second half of the first year of life (Lebedeva & Kuhl, 2010; Thiessen & Saffran, 2009). Rhythmic motor abilities are thought to provide relevant practice of rhythmic, closely timed actions, as they are required for speech (Iverson, 2010; Iverson et al., 2007). Yet, it remains unknown whether infants’ physical and neural tracking of music is also related to their language development.

Much of the research examining infant musicality, as reviewed above, used pre-recorded or digitalized stimuli that are easier to control and manipulate. However, pre-recorded music has a phenomenologically different quality compared with live musical performances (Trehub, 2017). Caregivers sing to infants on a daily basis (Steinberg et al., 2021), and ID singing is part of intimate interactions between caregivers and infants with the goal of engaging or soothing (Trainor et al., 1997). ID singing, compared to non-ID singing, is characterized by high pitch, slow tempo, enhanced temporal regularity, and greater dynamic range (Nakata & Trehub, 2011; Trainor et al., 1997). ID songs are often accompanied by idiosyncratic gestures (Eckerdal & Merker, 2009), energetic movement, and caregivers’ positive affect (Cirelli et al., 2020; Trehub et al., 2016), all of which are used by caregivers to attract infants’ attention, activate, and engage with them (Cirelli et al., 2020; Trehub & Trainor, 1998). Moreover, caregivers adjust their facial expressions and gaze to the timing of the ID songs (Lense et al., 2022). Still, it remains unclear how infants perceive and act upon *live* and *dynamic* ID singing.

The acoustic features of ID songs can be further specified by the context in which they are used, namely, to play or to soothe. Playsongs are characterized by higher rhythmicity, tempo, and pitch level, as well as more dynamic variability and thus increased complexity in comparison to lullabies (Cirelli et al., 2020; Rock et al., 1999; Trehub & Trainor, 1998). On the other hand, lullabies are characterized by lower tempo, pitch, and less variability, intending to soothe and calm an infant. Accordingly, the different acoustic features might affect how infants coordinate their neural and movement responses. Recent research in adults shows that neural tracking was enhanced in slower music (Weineck et al., 2022) and was guided by melodic expectations (Di Liberto et al., 2020). We thus expected enhanced neural tracking in slower and “simpler” ID songs (i.e., lullabies), while we hypothesized that the complexity of playsongs might attenuate infants’ neural tracking. Importantly, the specific acoustic features of lullabies and playsongs might have different effects on infants’ rhythmic movement than on neural tracking. On the behavioral level, increased rhythmic complexity was related to more movement and reported groove (i.e., the inclination to move) in children and adults (Cameron et al., 2022; Kragness et al., 2022). The enhanced complexity of playsongs might thus be more engaging to children.

Overall, we still know very little about infants’ neural tracking of and rhythmic movement to live and dynamic maternal singing. Closing this gap will help us understand how we might be able to facilitate infants’ physical, social, and cognitive development through musical communication. In the present study, we used the naturally occurring variance in acoustic features of playsongs and lullabies during live and dynamic ID singing to gain a deeper understanding of the tracking effects on infants’ bodies and brains. In Study 1, we measured 7-month-old infants’ neural tracking (using EEG) to maternal singing. To allow for more naturalistic and potentially more expressive movements, we observed rhythmic movements in an additional sample of 7-month-old infants while mothers sang to them (Study 2). We hypothesized that infants’ neural responses are more coordinated with lullabies than playsongs. In contrast, their rhythmic movements were expected to be more frequent with playsongs than lullabies. At 20 months, we assessed infants’ vocabulary to examine whether early musical engagement relates to infants’ language development. We hypothesized that higher neural tracking of ID songs, as well as more rhythmic movements, predict children’s vocabulary.

## Methods

### Participants

In Study 1, 30 7-month-old infants (13 females; age: 228.80 ± 8.20 days [*M* ± *SD*]) and their mothers (age: 33.96 ± 4.88 years) were included in the final sample. An additional 29 infants were tested but could not be included due to fussiness (*n* = 4), technical problems during the recording (*n* = 9), and bad signal quality due to the naturalistic setup (*n* = 16). In Study 2, 40 7-month-old infants (19 females; age: 227.17 ± 5.8 days) and their mothers (age: 35.10 ± 4.00 years) were included in the final sample. We excluded 7 additional mother-infant dyads from the final analysis due to infant fussiness (*n* = 5), or because the mother did not follow instructions (*n* = 2).

Participants were recruited from a database of families who expressed interest in participating in developmental research and have consented to being contacted by our staff. These families were recruited in neonatal units at local hospitals, in mother-child activity classes, and through social media. All infants were born full-term (38-42 weeks gestational age), had a birth weight of > 2500 g, and had no known developmental delays or neurological or hearing impairments. Infants grew up in predominantly German-speaking households. Mothers were highly educated, with 93.1% (Study 1) and 86.7% (Study 2) of mothers holding a university degree. Fifty-five percent of the mothers in both samples reported playing an instrument, and 20% were singing in a choir or a band. The study was conducted according to the declaration of Helsinki, approved by the university’s ethics committee, and parents provided written informed consent.

### Procedure

After arrival in the laboratory, parents and infants were familiarized with the environment. Parents were informed about the study and signed a consent form. In Study 1, the EEG setup was prepared while the infant sat on the parent’s lap. Infants were seated in an infant car seat, and their mothers faced them (*Figure 1A*). Study 2 followed the same procedure as Study 1, except infants sat upright in an infant highchair that allowed more movement. In Study 2, we additionally measured infants’ electro-cardiac rhythms using a two-lead derivation. These results will be reported elsewhere.

**Figure 1.**
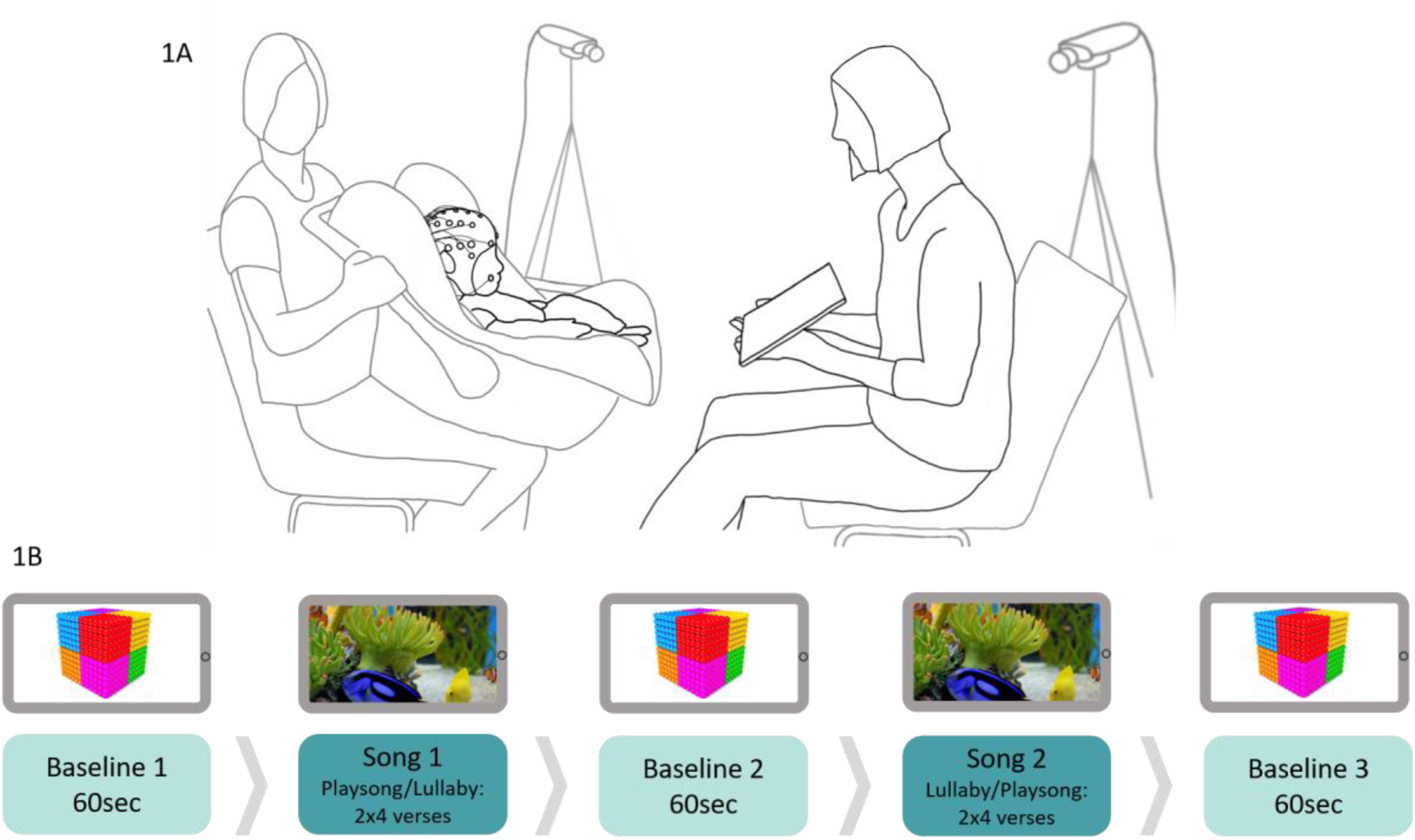
(A) Illustrated experimental setup. An experimenter held the car seat with the infant facing their mother in Study 1. Video cameras were used to record the mother-infant dyad from two angles. (B) Procedure. Mothers sang two repetitions of four verses for each lullaby and playsong. During both conditions, a tablet held by the mother played a video of fish swimming in an aquarium. A 60-second baseline, during which the mother did not interact with the infant, preceded, and followed each song, during which the tablet showed moving geometric shapes.

Each dyad was observed during two experimental singing blocks (*Figure 1B*), the order of which was randomized between participants. Each singing block was preceded and followed by a baseline (i.e., three in total), during which infants and mothers watched videos of slowly moving shapes for 60 s together. Mothers were instructed to refrain from talking during the baseline, but to reciprocate their infants’ communicative attempts (e.g., smiling, pointing towards the screen). During each singing block, mothers continued holding the tablet showing a calm aquarium video to help with infant fussiness. Mothers were instructed to sing one of two types of songs: a playsong or a lullaby. Songs were selected according to their level of familiarity during pilot testing. One example of each type of song was used. These were ‘Schlaf Kindlein, schlaf’ (lullaby) and ‘Es tanzt ein Bibabutzemann’ (playsong) (see SM Figure S1). All mothers sang the same songs and were informed about and provided recordings of the songs at the time of recruitment. Most (87%) of the mothers were familiar with the playsong, and 97% were familiar with the lullaby. All mothers sang four verses of each song and repeated each song, resulting in eight verses for each song. Mothers were prompted with a metronome before the commencement of each song (i.e., playsong = 170 bpm; lullaby = 100 bpm). Three cameras (synchronized in VideoSyncPro by Mangold International) were used to record the experiment.

### Measures

#### Maternal Singing

Audio of maternal singing was recorded using a portable microphone (Mangold International) at a 44,100 Hz sampling rate. Audio data were pre-processed using the software Audacity. <u>Excerpts</u> containing infant <u>vocalizations and</u> other noises from the environment were manually removed. To examine whether mothers differentiated between the two singing conditions, we analyzed maternal audio data in terms of mean and standard deviation in tempo (beats per minute from the autocorrelation function of the onset detection curve), pitch (Hz), root mean square (RMS; perceived loudness), and pulse clarity using MIRtoolbox (Lartillot et al., 2008). We also manually checked the derived tempo values by tapping along to the original recordings (random 10% of the sample) and found that our tapping values deviated from the tempo derived from MIRtoolbox by max. 2 BPM.

In order to extract the sound envelopes for further EEG analyses (Jessen et al., 2021), the audio data were then submitted to the NSL toolbox (http://nsl.isr.umd.edu/downloads.html). The resulting matrix contained band-specific envelopes of 128 frequency bands of uniform width on a logarithmic scale with center frequencies logarithmically spaced between 0.1 and 4 kHz. We obtained the broadband temporal envelope of the audio soundtrack by summing up band-specific envelopes across all frequencies to obtain one temporal envelope. The spectrogram of the envelope is included in the supplements (Figure S1). Lastly, the amplitude envelope data were down sampled to match the 500 Hz sampling rate of the EEG data.

#### Infant EEG (Study 1)

We recorded infant EEG using a BrainAmp DC 64-channel system and a 32-channel ActiCap system (Brain Products, Germany) containing active Ag/AgCl electrodes, mounted according to scalp locations of the 10-20 system. Horizontal and vertical electrooculograms were recorded bipolarly to control for eye movements. EEG data were sampled at 500 Hz. Impedances were controlled at the beginning of the experiment and accepted when below 20 kΩ.

EEG data were processed using the *eeglab* (Delorme & Makeig, 2004) and *fieldtrip* toolboxes (Oostenveld et al., 2010) as well as custom MATLAB code. EEG data were band-pass filtered with a noncausal finite impulse response filter (0.3 Hz - 100 Hz, −6 dB cutoff: 0.15 Hz - 100.15 Hz). Next, we reduced line noise (50/100 Hz) by using ZapLine from NoiseTools (de Cheveigné, 2020). Noisy channels were identified by assessing the normed joint probability of the average log power from 1 – 100 Hz and rejected if exceeding a threshold of 3 SD from the mean (number of removed channels: *M* = 4.61; *range* = 0 – 10). This step was repeated twice. Next, we used wavelet thresholding to identify and remove artifacts (threshold = hard, wavelet = coif4, level-dependent thresholding; Lopez et al., 2022). The previously rejected channels were then interpolated using spherical splines. Afterward, all channels were re-referenced to the average of all scalp electrodes (F7, F8, F9, F3, Fz, F4, FT7, FC3, FCz, FC4, FT8, T8 C4, Cz, C3, T7, CP3, CP4, TP10, P8, P4, Pz, P3, P7, PO9, PO10, O1, Oz, O2) and segmented into non-overlapping 1-s-epochs. In preparation for further analysis, epochs were automatically examined to see if they were still contaminated by artifacts. The standard deviation was computed in a sliding window of 200 ms. If the standard deviation exceeded 100 μV at any electrode, the entire epoch was discarded. Participants were excluded from further analysis if less than 90 artifact-free epochs could be retained. Infants’ processed EEG data comprised on average 144 epochs from the playsong condition and 126 epochs from the lullaby condition. The spectrogram of the data is provided in Figure S2. EEG data were then FIR-type bandpass filtered between 1 – 10 Hz (Jessen et al., 2019).

##### Neural tracking analyses

To quantify the degree to which the 7-month-old infants showed neural tracking of maternal singing, we used encoding models (i.e., temporal response functions [TRF]) to predict the infants’ EEG from the amplitude envelope of mothers’ singing. The TRF regresses the neural response onto the auditory signal, dividing out the autocovariance structure of the stimulus from the model. Further information on the approach is provided in the supplements.

Following Jessen et al. (2019), we estimated the predictive accuracy of the model by using individual models (computation of individual response function for each infant). We assumed individual models to be suitable for our purposes as ID songs were highly different between dyads, therefore leading to vastly different regressor weights between infants. To that end, 80% of the available data for a given participant was used to train the model. The resulting response function was then correlated with the response observed in the remaining 20% of the data. To accommodate a time lag between changes in the EEG signal and changes in the stimulus signal, predictions were computed over a range of time lags (i.e., 2 ms) between −100 and 500 ms later than the stimulus signal (see Jessen et al., 2021, for further details).

We obtained the optimal regularization parameter λ by training the respective model on the concatenated EEG and audio data of each infant using a variety of λ values between 10^-7^ and 10^7^. We increased the exponent in steps of 1 and used the resulting models to predict the EEG signal for each participant. By doing so, we obtained a total of fourteen different models (and predictions) based on the different λ parameters. For each of these 14 different models, we computed the mean response function across n-1 participants and used this response function to predict the EEG of the n^th^ fold (i.e., n-fold leave-one-out cross-validation). One fold comprised 1/5 of the data. Finally, we computed the predictive accuracy (i.e., Pearson’s correlation coefficient *r* between the predicted EEG and the actual EEG) for each infant and each electrode, resulting in 31 accuracy values per infant and stimulus parameters for each λ value. For each infant, stimulus parameter, and electrode set, we chose the λ value for which the model yielded the highest correlation between the predicted and the actual EEG time-series (Jessen et al., 2021).

For statistical evaluation, we computed two different predictive accuracies per infant and song conditions. First, we computed the correlation between the predicted response generated on a model trained on 80% of the data on the other 20% of the data (“individual model”). Second, a permuted or null predictive accuracy (“circular shift control”) was obtained. Before calculating accuracy this way, we reversed and applied a random circular shift to the true EEG time series for participant *n* (to ensure exceeding the potential autoregressive structure of the EEG) and computed the correlation between the shifted EEG and the predicted response from the individual model (for *n* = 400 permutations). The permutations were then averaged, resulting in one so-called “randomized” predictive accuracy value per infant.

To evaluate the resulting response functions, we computed a cluster-based permutation test with 1000 randomizations, testing the obtained response functions against zero. A cluster was defined along with the dimensions of time (lags between the sound envelope and the EEG) and electrode position, with the constraint that a cluster had to extend over at least two adjacent electrodes. A type-1 error probability of less than 0.05 was ensured at the cluster level (Meyer et al., 2021). The cluster-based permutation analysis revealed no significant clusters in the TRF, potentially due to the large individual differences between the infants’ evoked responses as well the acoustic features of the ID songs. The detailed results of this evaluation are reported in the supplements (see Figure S3).

#### Infant Rhythmic Movement

We coded the duration of infant rhythmic movement for both studies during the two singing conditions. Infant rhythmic movement was defined as at least three repetitions of the same movement in a body part or the entire body at regular short intervals (1 second or less; Thelen, 1979). We calculated proportionate movement values to compensate for differences in recording length. To ensure inter-rater reliability, three independent observers coded 30% of all videos yielding high inter-rater reliability with κ = .82.

#### Infant Gaze

To control for infant visual attention in both studies, we coded the following categories of infant gaze from video recordings using INTERACT by Mangold: (1) *gaze at mother’s face*; (2) *gaze at mother’s body; (3) gaze at tablet*; (4) *gaze away* as infant gaze directed away from their mother and the tablet at something else in their surrounding (Markova & Legerstee, 2006). Coding was conducted without sound, frame-by-frame, and each dwell time had to last at least 1 s. Inter-rater reliability was assessed on 30% of data coded by three pairs of independent observers, yielding moderate to excellent inter-rater reliability within a range of κ = .70-.99 for all categories. Analyses regarding infant gaze are reported in the supplements (Table S1).

#### Questionnaires

To assess language development, we used the Austrian Communicative Development Inventory - Level 2SF (ACDI-2 SF; Marschik et al., 2007) that mothers filled out when children were 20 months of age (age: 20.23 ±1.35 months [*M* ± *SD*]). Further self-reports on mothers’ depression symptoms, anxiety levels after singing, and familiarity of infants with the songs are included in the supplements.

### Statistical Analysis

All statistical analyses were conducted in RStudio (RStudio Team, 2022).

First, we ran a linear mixed-effects model comparing the prediction accuracy values between song type (playsong vs lullabies), data type (true (i.e., the predicted infant EEG from original sound envelopes correlated with the original infant EEG) vs shifted (i.e., the predicted infant EEG from the circular shifted sound envelope correlated with the original infant EEG)) in each channel. The prediction accuracy values were Fisher’s z transformed. The robustness of effects was evaluated using 95% confidence intervals and whether those excluded 0.

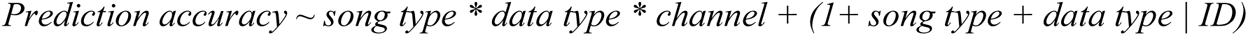

Next, we included the acoustic features of the playsong and lullaby (i.e., tempo, pitch, pulse clarity, and RMS) as fixed and interaction effects to examine whether differences in these features were related to differences in infants’ neural tracking of ID singing.

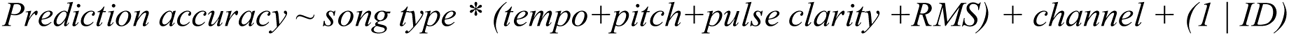

*Prediction accuracy ∼ song type * (tempo.sd+pitch.sd+pulse clarity.sd +RMS.sd) + channel + (1 | ID)*Finally, we included infants’ expressive vocabularies at 20 months as fixed and interaction effects to examine whether variation in their vocabulary was related to differences in infants’ neural tracking of ID singing.

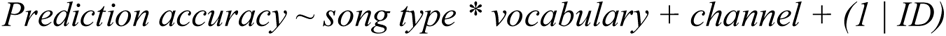

Next, we ran a generalized linear mixed-effects model assuming a beta distribution comparing the proportionate duration of rhythmic movements in the lullaby and playsong conditions in both studies.

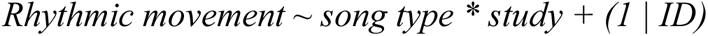

As we found significant differences in amounts of rhythmic movement between studies, we continued tested further relations with neural tracking and acoustic features separately for each study.

First, we tested whether neural coordination to ID songs in infants was related to the amount of rhythmic movement in Study 1.

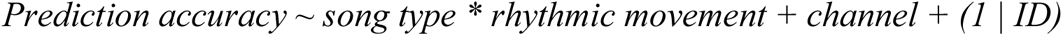

Next, we included the acoustic features of the playsong and lullaby (i.e., tempo, pitch, pulse clarity, and RMS) as fixed and interaction effects to examine whether differences in these features were related to differences in infants’ rhythmic movement during ID singing in Study 2.

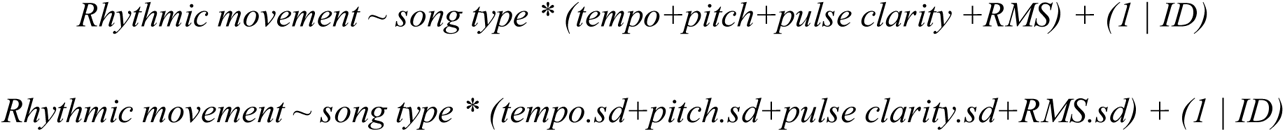

We included infants’ expressive vocabularies at 20 months as fixed and interaction effects to examine whether variation in their vocabulary was related to differences in infants’ neural tracking of ID singing during Study 1.

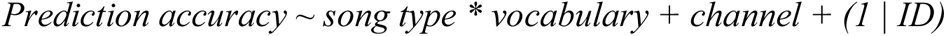

Finally, we included infants’ expressive vocabularies at 20 months as fixed and interaction effects to examine whether variation in their vocabulary was related to differences in infants’ rhythmic movement during ID singing during Study 2.

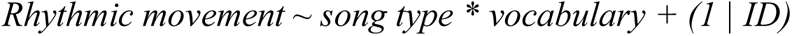

## Results

### Acoustic Features of Maternal Singing (Study 1 & 2)

First, we compared the acoustic features of the playsong and the lullaby (see Table 1 for descriptive statistics). Multiple comparisons were corrected for using the false discovery rate, which is indicated by the *q-*value (Benjamini & Hochberg, 1995). Playsongs were sung louder (χ^2^(1) = 34.15, *q* < .001), faster (χ^2^(1) = 40.77, *q* < .001), and in a higher pitch (χ^2^(1) = 8.83, *q* = .004) than lullabies. Playsongs and lullabies did not differ in pulse clarity (*q* = .248). Playsong varied more in perceived loudness (χ^2^(1) = 35.97, *q* < .001), tempo (χ^2^(1) = 6.79, *q* = .012), and pitch (χ^2^(1) = 57.50, *q* < .001) compared to lullabies. Lullabies varied more in pulse clarity compared to playsongs (χ^2^(1) = 4.505, *q* = .034).

**Table 1.**
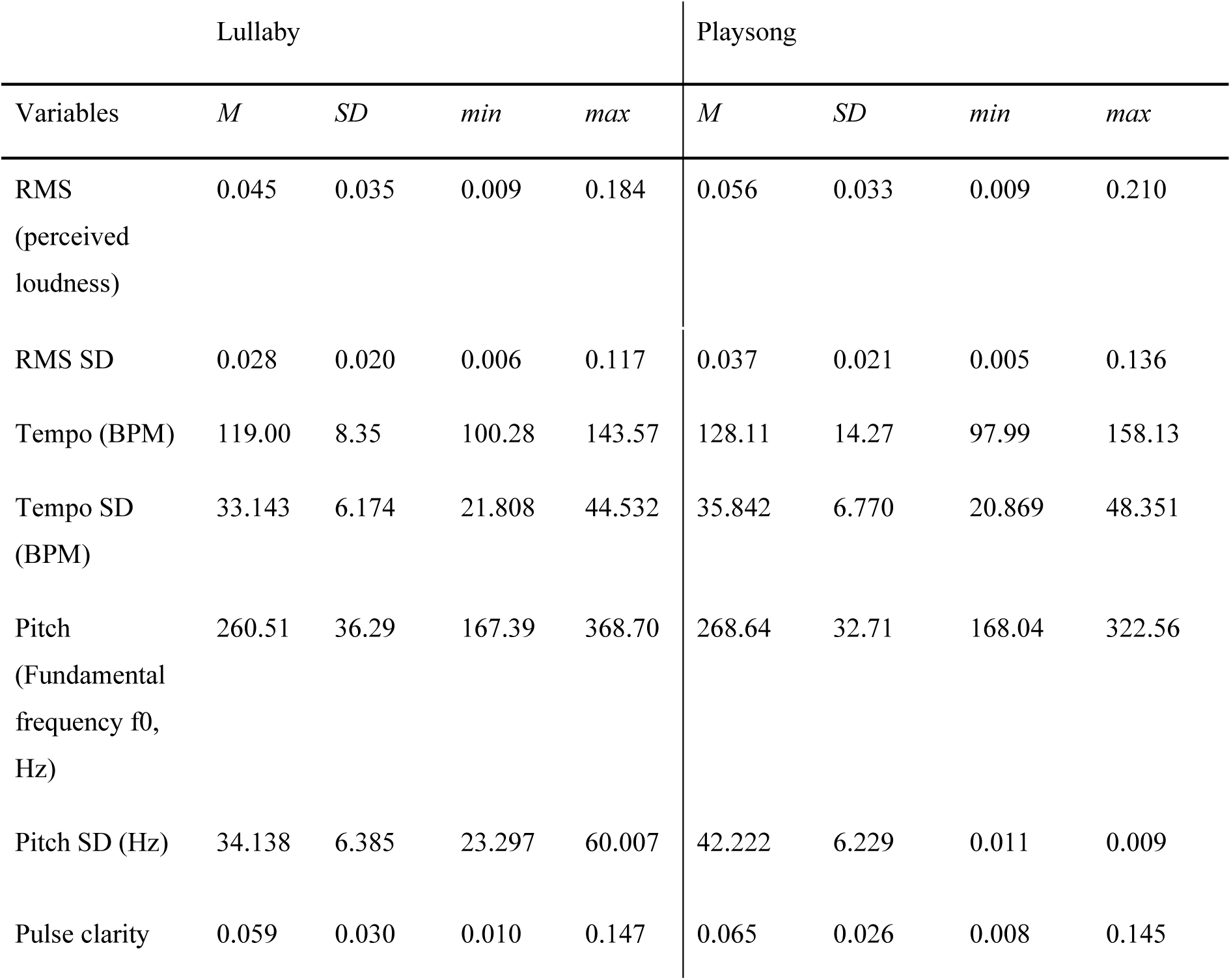
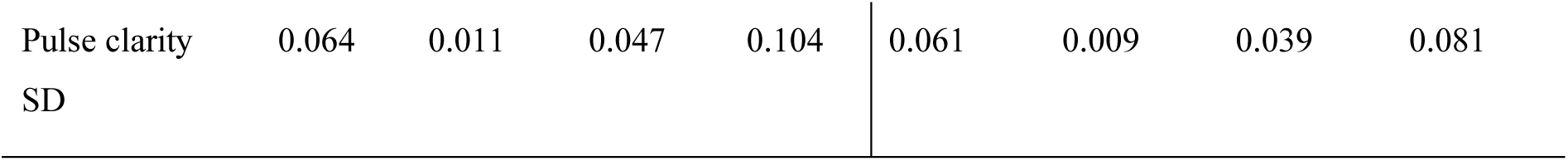
Descriptive statistics on acoustic features of live-sung lullabies and playsongs (Study 1 & 2)

### Neural Tracking of Maternal Singing (Study 1)

First, we compared the predictive accuracy obtained by the individual models of both playsong and lullaby. The model results displayed a significant fixed effect of ID song type (χ^2^(1) = 42.135, *p* < .001), a fixed effect of data type (χ^2^(1) = 2430.361, *p* < .001), and an interaction effect between ID song type and data type, χ^2^(1) = 40.881, *p* < .001. The set of effects was further tested in post-hoc contrasts (corrected for multiple comparisons) and showed that infants’ neural tracking of ID singing (true pairing) was more accurate (estimates = 0.065, SE = 0.002, 95% CI = [0.062 0.068]) in comparison to the randomized control (shifted pairing; estimates = 0.001, SE = 0.002, 95% CI = [0.004 0.004]). Thus, infants showed significant neural tracking of both lullabies and playsongs. In addition, neural tracking of lullabies (estimates = 0.074, SE = 0.002, 95% CI = [0.070 0.077]) was more accurate than that of playsongs (estimates = 0.057, SE = 0.002, 95% CI = [0.053 0.060], Figure 2A) in data type true, while neural tracking of the two song types did not differ in data type shifted, *p* = .999. Other effects, such as the fixed effect of channel, the interaction effect between channel and type of song, the interaction effect between channel and data type, and the three-way interaction between channel, type of song, and data type were not significant (*p* > .086). In sum, we found above-threshold neural tracking of lullabies and playsongs, while this was more accurate for lullabies than playsongs across the infant brain.

**Figure 2.**
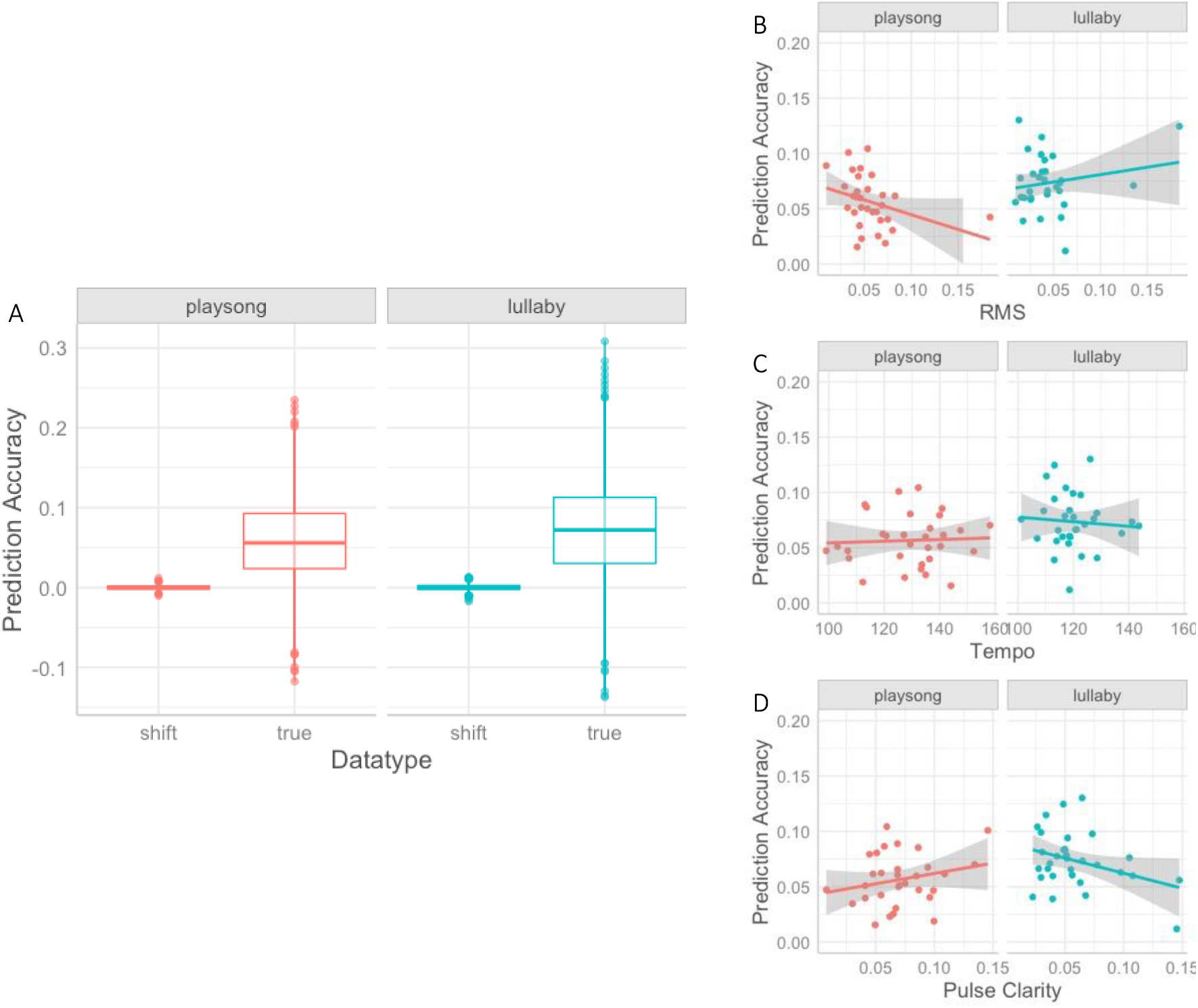
(A) Graph depicting the interaction effect between true vs. shifted data (x Axis) in the different types of ID songs (columns) on prediction accuracy (y Axis). The difference in prediction accuracy (y Axis) between song types (columns) is modulated by (B) the perceived loudness of playsongs (x Axis), (C) the tempo of lullabies (x Axis), and (D) the pulse clarity of playsongs (x Axis).

#### Neural tracking and acoustic features

Next, we examined the effects of tempo, pitch, pulse clarity, RMS on the predictive accuracies of TRF models tested on the original data. First, the mean of acoustic features was included as fixed and interaction effects in the linear mixed-effects model. The model showed that the fixed effect of ID song type remained when controlling for the acoustic features, χ^2^(1) = 16.164, *p* < .001. There were, however, significant interaction effects between ID song type and RMS, χ^2^(1) = 14.264, *p* < .001 (Figure 2B), ID song type and tempo, χ^2^(1) = 4.925, *p* = .026 (Figure 2C), as well as ID song type and pulse clarity, χ^2^(1) = 9.796, *p* = .002 (Figure 2D). Post-hoc analyses of the interaction effects showed that lower RMS (perceived loudness) was related to more accurate neural tracking of the playsong (*trend* = 0.015, *SE* = 0.004, *95% CI* = [-0.024 −0.008]), while RMS was not related to neural tracking accuracy of lullabies (*trend* = −0.003, *SE* = 0.003, *95% CI* = [-0.009 0.002]). Moreover, slower-sung lullabies were related to more accurate neural tracking (*trend* = −0.008, *SE* = 0.004, *95% CI* = [-0.016 −0.001]), while the tempo of playsongs did not relate to neural tracking (*trend* = 0.000, *SE* = 0.002, *95% CI* = [-0.004 0.005]). Higher pulse clarity in playsongs was further related to enhanced neural tracking (*trend* = 0.010, *SE* = 0.003, *95% CI* = [0.004 0.015]) but did not affect neural tracking of lullabies (*trend* = −0.003, *SE* = 0.002, *95% CI* = [-0.008 0.002]).

#### Neural tracking and acoustic variability

In the next step, we tested the variability of the acoustic features as fixed and interaction effects on neural tracking. We found significant fixed effects of ID song type (χ^2^(1) = 8.866, *p* = .003) and variability of tempo (χ^2^(1) = 5.972, *p* = .015), indicating that neural tracking was enhanced when the tempo of ID songs varied less, *estimate* = −0.008, *SE* = 0.003, *95% CI* = [-0.013 −0.002]. There were also significant interaction effects between ID song type and RMS variability, χ^2^(1) = 11.762, *p* = .001, as well as ID song type and pitch variability, χ^2^(1) = 11.329, *p* = .008. Post-hoc analyses showed that lower RMS variability was related to more accurate neural tracking of playsongs (*trend* = −0.011, *SE* = 0.004, *95% CI* = [-0.019 −0.004]), while RMS variability was not related to neural tracking of lullabies (*trend* = −0.000, *SE* = 0.003, *95% CI* = [-0.006 0.005]). Lower pitch variability was related to enhanced neural tracking of lullabies (*trend* = −0.006, *SE* = 0.003, *95% CI* = [-0.012 −0.001]), while it was not significantly associated with neural tracking of playsongs (*trend* = 0.006, *SE* = 0.004, *95% CI* = [-0.001 0.013]). All other acoustic features were not related to predictive accuracy (*p* > .063).

### Rhythmic Movement to Maternal Singing (Study 1 & 2)

Based on planned analysis, we conducted rhythmic movement analyses separately for both studies. The overall model, including both studies together, is reported in the Supplements.

#### Effects of song type on rhythmic movements (Study 1)

Generalized linear models reveal that rhythmic movement did not differ significantly between song types in Study 1 (*z*=1.418, *p*=.156; Figure 3, left panel).

**Figure 3.**
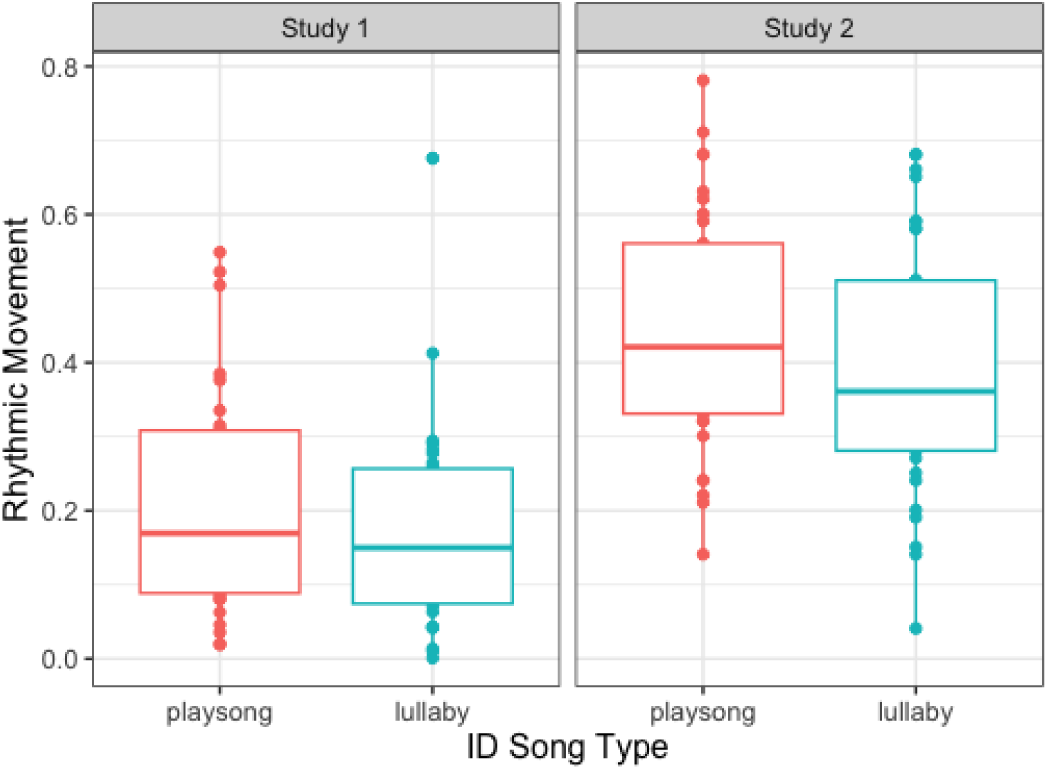
Graph depicting the fixed effect of the type of ID song (x Axis) on rhythmic movement frequency proportionate to the condition duration (y Axis) in Study 1 (left panel) and Study 2 (right panel).

#### Neural tracking and rhythmic movement (Study 1)

We tested whether rhythmic movements in infants were related to neural tracking of maternal singing. Here, the model outputs showed a significant fixed effect of rhythmic movement, χ^2^(1) = 10.816, *p* = .001, indicating that more frequent rhythmic movements were related to more accurate neural tracking of maternal singing, independent of song type, *estimate* = 0.027, *SE* = 0.011, *95% CI* = [0.007 0.049]. Importantly, the fixed effect of song types remained, χ^2^(1) = 34.594, *p* < .001 (lullabies were better tracked than playsongs), while the interaction effect between rhythmic movement and song type was not significant, *p* = .941.

#### Effects of song type on rhythmic movements (Study 2)

Next, we ran the planned and more detailed rhythmic movement analysis (including acoustic features and language) for Study 2. Infants’ rhythmic movement differed significantly between song types in Study 2 (*z*=2.172, *p*=.029, Figure 3, right panel).

#### Acoustic features and rhythmic movement (Study 2)

Next, we tested the role of the acoustic features of maternal ID singing on infants’ rhythmic movement. We found no significant association between mean values and standard deviations of acoustic features and the amount of rhythmic movement (*p* > .070).

### Relations Between Neural Tracking (Study 1), Rhythmic Movement (Study 2), and Children’s Language Development

Finally, we examined whether infants’ neural tracking and rhythmic movement in response to maternal ID singing are related to their expressive vocabulary at 20 months. The model outputs for Study 1 (N=27) revealed a significant interaction effect of neural tracking and ID song type on vocabulary size at 20 months, χ^2^(1) = 5.236, *p* = .022. The post-hoc analysis (see Figure 4A) revealed a significant, but not robust, trend for enhanced neural tracking of playsongs in relation to larger vocabulary size at 20 months (*trend* = 0.002, *SE* = 0.002, *95% CI* = [-0.002 0.006]), and no significant relation between neural tracking of lullabies and vocabulary size (*trend* = −0.002, *SE* = 0.002, *95% CI* = [-0.006 0.002]). Results for Study 2 (N=33) showed that infants’ rhythmic movement was positively related to infants’ expressive vocabulary depending on the song type, χ^2^(1) = 4.933, *p* = .026.

**Figure 4.**
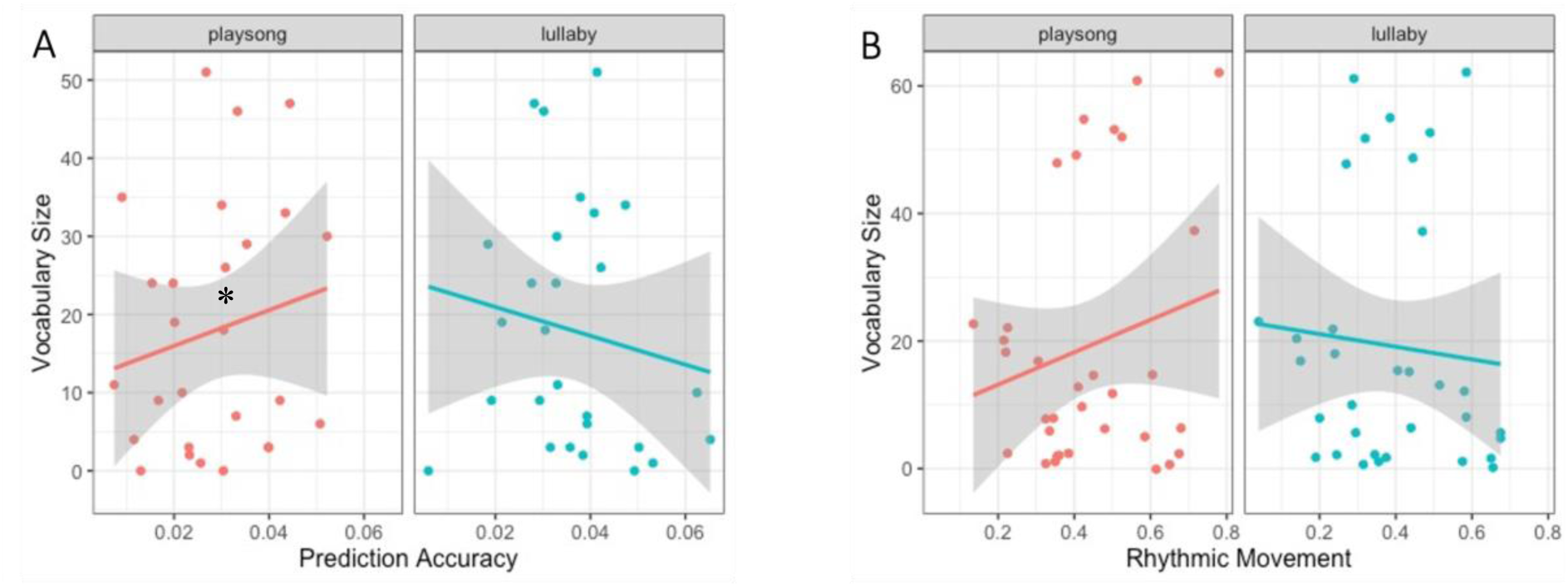
(A) Graph depicting the relation between neural tracking (x Axis) and (B) rhythmic movement at 7 months (x Axis) in playsongs (red) and lullaby (blue) on estimated means of vocabulary size at 20 months (y Axis). Shaded area indicates the 95% confidence interval.

Specifically, post-hoc trend analysis showed that infants’ rhythmic movement during the playsong, but not the lullaby, was positively related to their vocabulary (see Figure 4B). The trend was, however, not robust (*trend* = 0.072, *SE* = 0.068, *95% CI* = [-0.062 0.205]).

## General Discussion

In the two studies reported here, we investigated 7-month-old infants’ neural tracking of and rhythmic movement to live and dynamic maternal ID singing. In addition, we tested whether and how these forms of coordination are related to children’s language development. We found that 7-month-old infants coordinate both at the neural and behavioral levels with their mothers’ live ID singing. In line with our hypotheses, infants showed better neural tracking of lullabies than playsongs (in Study 1), while they displayed more rhythmic movements during playsongs than during lullabies (in Study 2). Interestingly, only infants’ neural and behavioral coordination with their mothers’ playsongs, but not lullabies, predicted their vocabulary size at 20 months.

Infants showed above-threshold neural tracking of live-sung ID songs in comparison to a shuffled control. This finding highlights that infants are able to neurally track live musical communication beyond rhythmic pure tones (Cirelli et al., 2016), recordings of nursery rhymes (Attaheri, Choisdealbha, Di Liberto, et al., 2022), or infant-directed speech (Menn, Michel, et al., 2022). Neural tracking reflects the degree to which brain activity regresses onto a continuous stimulus (such as volume fluctuations in song; e.g., Jessen et al., 2019; Kalashnikova et al., 2018). Specifically, our encoding approach, based on TRF, quantifies the degree to which infant ongoing brain activity can be “predicted” by the sound envelope of maternal singing of a lullaby vs. playsong. The closer the infant brain tracks the stimulus, the higher the derived encoding accuracy, that is, the better we can estimate the individual baby’s brain response based on the sound envelope of their mother’s singing. In adults, directing top-down selective attention towards a stimulus associates with increased neural tracking (e.g., Zion Golumbic et al., 2013). These results indicate that infants’ neural tracking of live-sung ID songs could reflect attentional processes as well. In addition, neural tracking depends on stimulus features that can facilitate or obstruct neural processing. For instance, maternal infant-directed speech contains amplitude modulations that make it particularly easy to track for infants (Menn, Michel, et al., 2022).

Interestingly, our results indicate that infants were more accurate at tracking lullabies than playsongs even when controlling for concurrent rhythmic movements. Lullabies are sung with a regular and repetitive pattern (Cirelli et al., 2020; Trainor et al., 1997) that could help infants to neurally track this type of song. This interpretation is supported by acoustic analyses of maternal singing in both studies showing that lullabies, compared to playsongs, were sung slower, lower in pitch, and less variable, possibly making them more predictable. These individual acoustic features of lullabies were further related to enhanced neural tracking, such that individual lullabies sung at a slower tempo also showed enhanced neural tracking (Figure 2C). This is consistent with previous findings with adults which show enhanced neural tracking of slower music (Weineck et al., 2022). Our results thus corroborate the relation between a clear and predictable structure in tempo and pitch and potential ease of neural coordination. This increased clarity in tempo- and pitch-related structures in lullabies might also be particularly associated with their regulatory functions, such as modulating emotions (Trehub et al., 2015) and maintaining infant attention and composure (Shenfield et al., 2003). Despite their more variable and faster structure, infants were also able to neurally track playsongs (Study 1) and actually showed more rhythmic movement during playsongs than lullabies (Study 2). In line with our argumentation, neural coordination was enhanced when caregivers sang with higher pulse clarity, quieter, and less variability in loudness. We, therefore, suggest that caregivers might counterbalance the complexity of playsongs by enhancing their rhythmic structure (Zentner & Eerola, 2010) as well as using loudness to guide infants’ attention (Calignano et al., 2021). In previous research, pulse clarity was also related to infants’ amounts of rhythmic movement (Zentner & Eerola, 2010). Generally, our results suggest that variability in acoustic features, potentially up to a certain threshold, does not hinder infants’ neural tracking of ID songs. Instead, increased rhythmic variability might increase engagement (Cameron et al., 2022). The discrepancy in rhythmic movement frequency between playsongs and lullabies is likely further pronounced by infants’ familiarity with the songs (see Supplements) and potentially their expectation to engage in an activating social context. Alternatively, rhythmic movement might arise to help infants process these acoustically more variable ID songs, as we found that more accurate neural tracking in infants was related to more rhythmic movements irrespective of song type. Even though in our studies we could not distinguish between musical and non-musical rhythmic behavior, previous research suggests that the perception of rhythm while listening to music often involves spontaneous movement on these periodicities (Hurley et al., 2014; Toiviainen et al., 2010). In adults, rhythmic motor activity facilitates processing of on-beat tones through temporal alignment of attention fluctuations (Morillon et al., 2014; Zalta et al., 2020). Music stimulates movement and activates motor pathways (Grahn & Brett, 2007; Vuilleumier & Trost, 2015), and already 5-month-olds move rhythmically significantly more to music than speech, suggesting that there may be a predisposition for rhythmic movement in response to music (Zentner & Eerola, 2010). In the current study, the types of rhythmic movements included were 1) whole-body movements where infants used the footstep of the highchair to move their body upwards, and 2) kicking/tapping their feet or flapping their arms and moving their fingers. Interestingly, passive body movement was also found to influence 7-month-old infants’ encoding of musical rhythms (Phillips-Silver & Trainor, 2005), indicating that their own motor experiences may contribute to young infants’ processing of more variable acoustic input.

Importantly, we found that infants’ diverging coordination to live and dynamic lullaby and playsong was also reflected in the association with language outcomes at 20 months. Results showed that infants’ neural tracking of and rhythmic movement to playsongs, but not lullabies, were positively related to their later vocabulary size, even though the relation was not robust. This finding supports the proposition that musicality evolved as a common ability for infants to decipher music and language (Brandt et al., 2012). Preverbal infants, who are not yet familiar with the rules of language, classify speech according to pitch, melody, and rhythm in order to understand its meaning (Goswami, 2012). It is precisely prosodic information (i.e., slow speech rhythms) that is crucial for language acquisition (Goswami, 2012; Langus et al., 2017). Neural tracking of the more variable playsong might, for example, relate to infants’ ability to extract prosodic information from a vocal stream and thus possibly support their word segmentation abilities (Menn, Ward, et al., 2022), marked by larger expressive vocabulary size at age 20 months. ID singing may be particularly helpful for language acquisition, in addition to ID speech, due to its metric structure that makes prosodic stress more pronounced (Nakata & Trehub, 2011; Suppanen et al., 2019; Trainor et al., 1997). In addition, moving rhythmically could help infants process the prosodic information from the song, a notion supported by arguments that rhythmic abilities are a central part of language acquisition (Thomson & Goswami, 2008). An alternative explanation we cannot rule out is that infants’ movements affected the EEG response; movement-related artifacts in the EEG might have increased infant neural tracking in the playsong and thus the relation between neural tracking and infants’ later language development. Nonetheless, our study provides promising evidence for the connection between early musical experiences and language development.

The paradigm has some limitations. Mothers’ singing was highly varied on the individual level, which did not allow us to make stronger claims about the relation of specific acoustic features (such as pitch and tempo) in relation to neural tracking. Hence, future studies might consider more controlled stimuli that vary less across infants (e.g., Lense et al., 2022; Nguyen et al., 2023; Weineck et al., 2022) or use methodological approaches that can take into account how the variance in stimuli relates to infants’ tracking/attention over time. In addition, we need to consider the fit of the musical piece to the particular situation in which we observed infants. The context of the laboratory setting likely afforded the playsong more than the lullaby, as the mothers and the experimenter intended to keep infants attentive and entertained. Thus, the congruence between the observational situation and the playsong may have contributed to our pattern of results. Moreover, it is possible that mothers themselves moved differently during the two types of songs, and infants simply tracked and imitated their mothers’ movement type and frequency. Playsongs, in particular, are not only expressed through voice but also the body and are often intrinsically multimodal (Eckerdal & Merker, 2009; Lense et al., 2022). In fact, our supplementary analyses showed that infants looked significantly longer at their mothers during the playsong than during the lullaby. However, infant gaze behavior during the playsong was neither associated with their neural tracking nor rhythmic movements in that condition. Instead, we found that neural tracking of lullabies was weaker when infants looked away from mothers’ faces and bodies, suggesting that neural tracking is related to infants’ attention towards the face (Lense et al., 2022). To fully disentangle the factors underlying infants’ neural tracking and rhythmic movements during ID songs, future studies need to include more in-depth analyses of the accompanying sensorimotor synchronization processes, including caregivers’ movements (e.g., rocking, playful gestures, etc.).

## Conclusion

Singing represents a flexible tool for caregivers to meet the various needs of their infants. However, it remains a question of great theoretical controversy why ID singing is such a powerful communication method with very young infants (Mehr et al., 2021; Savage et al., 2021). Markova et al. (2019, 2020) argued that ID singing has inherent acoustic characteristics that support infants’ coordination to a particular interactive rhythm purported by the caregiver, and this process could have important developmental implications. Findings of the present study corroborate these hypotheses by showing that lullabies may be used to help infants track a slower and regular rhythmic pattern, while playsongs, containing more variability, could be perceived as more engaging and thus make infants move rhythmically.

The acoustic variability of playsongs also seems to promote language acquisition in that rhythmic variability may draw attention to the songs’ prosodic information (Hannon & Johnson, 2005). These findings highlight potential avenues for future studies to test the neural and behavioral basis of the communicative function of mother-infant musical interaction by experimentally manipulating the relevant acoustic features. With its semi-naturalistic design, the present study provides a better understanding of the dynamics, mechanisms, and developmental outcomes of early musical interactions.

## Acknowledgments

We are grateful to Liesbeth Forsthuber, Christina Schaetz, Martina de Eccher, Sarah Paul, Marlies Schermann, Anna Matyk, Rahel Loos, Madlene Radosavljevic, Pierre Labendzki, Jakob Weickmann and Elisa Roberti for their support in data acquisition, processing, and video coding. Additionally, we thank all families who participated in the study and the Department of Obstetrics and Gynecology of the Vienna General Hospital for supporting our participant recruitment.

## CRediT author statement

*Trinh Nguyen*: Conceptualization, Methodology, Formal analysis, Data curation, Writing – Original Draft, Writing – Review & Editing, Visualization, Project administration *Susanne Reisner*: Data curation, Writing – Original Draft, Writing – Review & Editing, Visualization *Anja Lueger*: Investigation, Data – Curation, Writing – Original Draft *Sam V. Wass*: Methodology, Formal analysis, Writing – Review & Editing *Stefanie Hoehl:* Conceptualization, Methodology, Resources, Writing – Review & Editing, Supervision, Funding acquisition *Gabriela Markova:* Conceptualization, Methodology, Investigation, Data curation, Writing – Original Draft, Writing – Review & Editing, Supervision, Project administration

