## Supplementary Information for "Sing to me, baby: Infants show neural tracking and rhythmic movements to live and dynamic maternal singing"

**Figure S1. Notations and lyrics of the lullaby (“Schlaf, Kindlein, schlaf”) and the playsong (“Es tanzt ein Bibabutzemann”).**

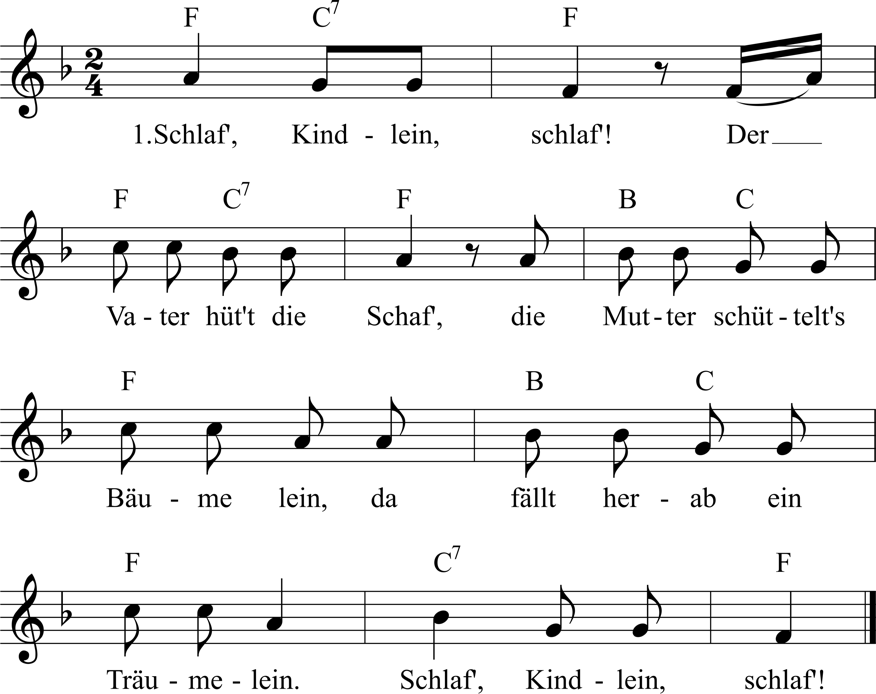

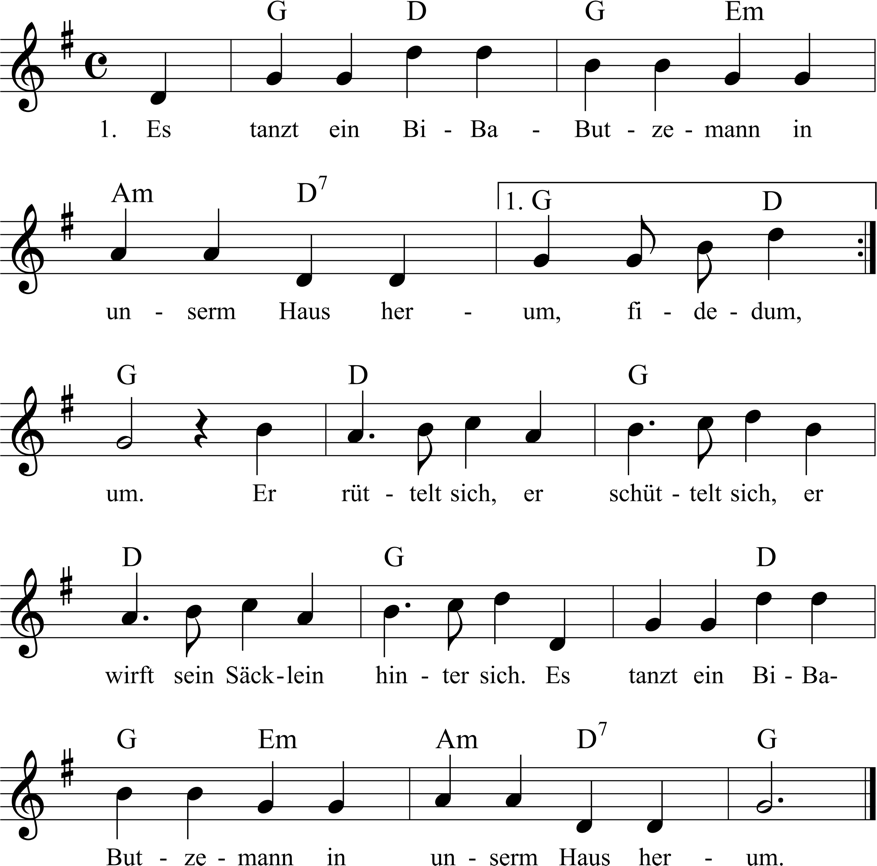

Note. The lullaby is a German children’s song in F-major, and we primed participants to sing the song at 100 BPM using a metronome. The playsong is a German children’s song in G-Major, and participants were primed to sing the song at 170 BPM. Each verse in the songs lasted on average 14.6 s (lullaby) to 22.3 s (playsong), and mothers repeated the verse eight times in total, amounting to average audio lengths of 116.42 s (lullaby; range = 96-207 s) and 178.25 s (playsong; range = 113-232 s).

**Maternal depression symptoms and anxiety levels after singing**

All participating mothers completed the Edinburgh Postnatal Depression Scale (EPDS; Bergant et al., 1998), to control for depressive mood (*M* = 4.83, *SD* = 3.64, *range* = 0-13). As mothers showed subrange (≥13) depression symptoms, we were able to include all dyads in subsequent analyses. Mothers also completed the State-Trait Anxiety Inventory (STAI; Marteau & Bekker, 1992) upon the completion of the experiment, which captured their low to medium anxiety levels after singing (*M* = 22.51, *SD* = 11.86, *range* = 1.43 - 47.13).

**Removal of background noise including infant vocalizations**

We manually removed excerpts containing environmental noises from the audio recording using the software Audacity (<https://www.audacityteam.org/>). These excerpts included noises the infant made with their body (e.g., hitting against the highchair), vegetative noises (e.g., coughs and burps), infant vocalizations, and other background noise. The removed contaminations did not differ significantly between the lullaby and playsong conditions, both in frequency (*V* = 201, *p* = .52) and relative duration (*V* = 225, *p* = .89). Additionally, infant vocalizations did not differ significantly in frequency (*V* = 63.5, *p* = .07) or relative duration (*V* = 102, *p* = .17) between the lullaby and playsong conditions. To ensure inter-rater reliability, two independent observers coded manually removed audio excerpts on whether they contained infant vocalizations in 30% of all audios, yielding high inter-reliability with κ = .93.

**Assessment of infant exposure to music**

We collected self-reported information on infant exposure to music: Families listened to music for an average of 14.38 hours/week (*SD* = 15.37, *range* = 2-70); mothers sang to infants for an average of 5.86 hours/week (*SD* = 4.79, *range* = 0-20); mothers made music to infants for an average of 3.72 hours/week (*SD* = 3.80, *range* = 0.5-21).

**Familiarity with ID songs**

We then tested whether infants’ familiarity with the songs was associated with how well they tracked or rhythmically moved to the ID songs. Familiarity (based on maternal self-report) was not significantly related to the ID songs, *p* > .057. However, familiarity was related to infants’ rhythmic movement to the different types of ID songs. Results revealed a significant interaction effect between the type of ID song and infants’ familiarity with the playsong, Χ^2^(1) = 7.407, *p* = .006. Post-hoc analysis revealed that infants' difference in rhythmic movement to the two types of ID songs was more pronounced when they were more familiar with the playsong, *contrast* *estimate* = 0.096, SE = 0.035, 95% CI = [0.027 0.165].

**Additional information on TRF**

The response model is defined as:

$r\left( t,n \right)= \sum_{l} w\left( l,n \right)s\left( t-l \right)+\varepsilon(t,n)$

where $r\left( t,n \right)$ is the neural response, across each channel included in the model *(n)*, across all time points *(t);* *w* is the TRF that describes the linear transformation of the stimulus *(s)* to the ongoing neural response *(t)*, and $\varepsilon$ is the residual error at each channel not explained by the model.

The TRF (w) is estimated using the following equation:

$$w={(S^{T}S+\lambda I)}^{-1}S^{T}r$$

where $S^{T}r$ is the result of convolution between the zero-lagged neural response *(r)*, and the speech stimulus at different time lags (S), $S^{T}S$ is the autocovariance matrix of the stimulus at each time lag which is divided out from the linear relationship modeled in $S^{T}r$. I is the identity matrix, and $\lambda$is the ridge parameter, which enforces a smoothness constraint on the output of the model (w). The output (w) is a time-windows by EEG channels matrix, containing the resulting TRF weights for each channel at each time window (see Crosse et al., 2016 for further explanations). A constant term is included in the model by concatenating a column of ones to the left of *S*.

A method of ridge regression is used to improve the reliability of the estimated coefficients and prevent over-fitting. Ridge regression works by introducing a bias term to the model, which penalizes TRF values as a function of their distance from 0 (Crosse et al., 2016). This reduces over-estimation problems in *w* and reduces the likelihood of high frequency artifacts in the estimated TRF model (Fiedler et al., 2017; Haufe et al., 2014) .

**Gaze behavior**

Table S1 shows infants’ gaze behaviors towards their mother, the tablet, and away from both as proportions according to the respective lengths of the songs. We tested whether infant gaze behavior differed between the two conditions.

Paired t-tests revealed that infants looked significantly longer at their mothers in the playsong condition in comparison to the lullaby condition, *t*(70) = 2.56, *p* = .013. Infants’ gaze at either the tablet or away did not significantly differ between conditions, *p* = .680, *p* = .452.

**Table S1.** Descriptive statistics of infant’s gaze behavior

|  | Lullaby | | | | Playsong | | | |
| --- | --- | --- | --- | --- | --- | --- | --- | --- |
| Variables | *M* | *SD* | *min* | *max* | *M* | *SD* | *min* | *max* |
| Gaze towards mother’s face (%) | 12.38 | 12.10 | 0.00 | 70.39 | 15.73 | 15.42 | 0.00 | 76.08 |
| Gaze towards mother’s body (%) | 0.08 | 0.40 | 0.00 | 2.40 | 0.05 | 0.37 | 0.00 | 3.02 |
| Gaze towards tablet (%) | 58.29 | 20.77 | 0.00 | 97.92 | 55.17 | 22.10 | 10.40 | 97.42 |
| Gaze away (%) | 28.08 | 18.22 | 0.00 | 70.10 | 28.31 | 18.68 | 0 | 89.60 |

**Gaze and neural tracking**

Next, we examined whether neural tracking in both conditions was associated with the infants’ gaze behaviors. We, therefore, separately included the infants’ gaze behaviors during the playsong and lullaby (i.e., gaze towards mother, gaze at the tablet, and gaze away) as fixed and interaction effects, to examine whether differences in these features were related to differences in infants’ neural tracking of ID singing. Infants, who looked away for longer durations during the lullaby condition, also showed lower predictive accuracy, namely weaker neural tracking, *estimate* = -0.038, SE = 0.018, 95% CI = [-0.074 -0.004], Χ^2^(1) = 11.94, *p* < .001. Infants' other-looking behavior was not related to how well they neurally tracked their mother’s singing, *p* > .090.

**Gaze and rhythmic movement**

We tested whether infants’ gaze behavior affected their rhythmic movement. We, however, found no significant associations between infant gaze and their rhythmic movement, *p* > .115.

**Trial duration and neural tracking**

Next, we analyzed whether infants’ neural tracking was associated with the length of the conditions. We, therefore, added trial durations as a fixed effect to the linear mixed effects model:

*Prediction accuracy ~ song type * trial duration + channel + (1 | ID)*

The model output revealed that trial duration was not significantly related to neural tracking, *p*=.604. While the interaction effect between song type and trial durations was significant (Χ^2^(1) = 5.85, *p* = .016, the estimated slopes were not robust (lullaby: emtrends = 0.002, SE = 0.002, 95% CI = [-0.001 0.004]; playsong: emtrends = -0.003, SE = 0.002, 95% CI = [-0.006 0.000].

**Trial duration and gaze behavior**

We also analyzed whether infants’ gaze behavior was associated with the length of the conditions. We, therefore, added trial durations as a fixed effect to the linear mixed effects model:

*Trial duration ~ song type * (gaze at mother + gaze at body + gaze away) + (1 | ID)*

The model output revealed that trial duration was not significantly related to infants’ gaze behavior to the mother, *p* = .959, but that trial duration was shorter when infants looked away for longer (Χ^2^(1) = 4.45, *p* = .035, *emtrends* = -51.812, SE = 28.832, 95% CI = -106.65 2.784]. The effect was, however, not robust.”

**Additional Rhythmic Movement Analyses**

We also assessed rhythmic movements in Study 1 and Study 2 together and tested whether infants showed more rhythmic movement during the lullaby or the playsong condition in both studies. The generalized linear mixed-effects model output displayed a significant effect of song type, χ^2^(1) = 6.474, *p* = .011. Model estimates revealed that infants showed less rhythmic movement in the lullaby condition (*estimate* = 0.250, *SE* = 0.020, *95% CI* = [0.212 0.292]) than in the playsong condition (*estimate* = 0.296, *SE* = 0.022, *95% CI* = [0.254 0.341]). Given the methodological differences between studies (i.e., that infants were restrained in Study 1 but not in Study 2), we also find significant differences between studies, χ^2^(1) = 40.893, *p* < .001. As predicted, infants moved more in Study 2 (*estimate* = 0.410, *SE* = 0.031, *95% CI* = [0.350 0.472] than in Study 1 (*estimate* = 0.168, *SE* = 0.020, *95% CI* = [0.132 0.211]). The interaction effect was not significant, *p*=.860.

**Frequency information of EEG and sound envelopes**

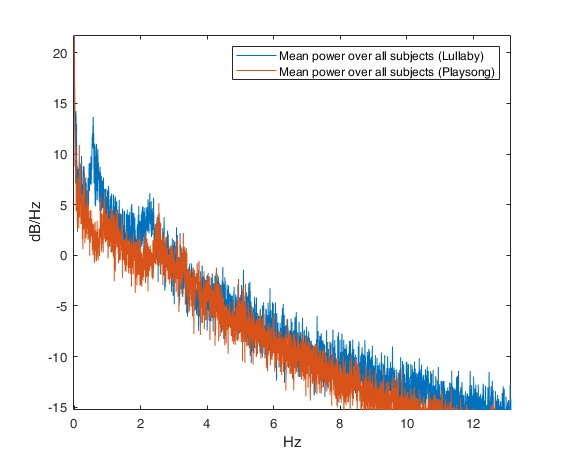

**Figure S2**. Frequency plot of amplitude-modulated sound envelopes for ID songs averaged over each condition (lullaby = blue, playsong = red). Frequency is plotted on the x Axis, and power in dB/Hz is on the y Axis. The songs show beat- (1.9-3.4 Hz) and meter-related (0.5-1.2 Hz) peaks relevant to the tempo of each condition in the spectrogram.

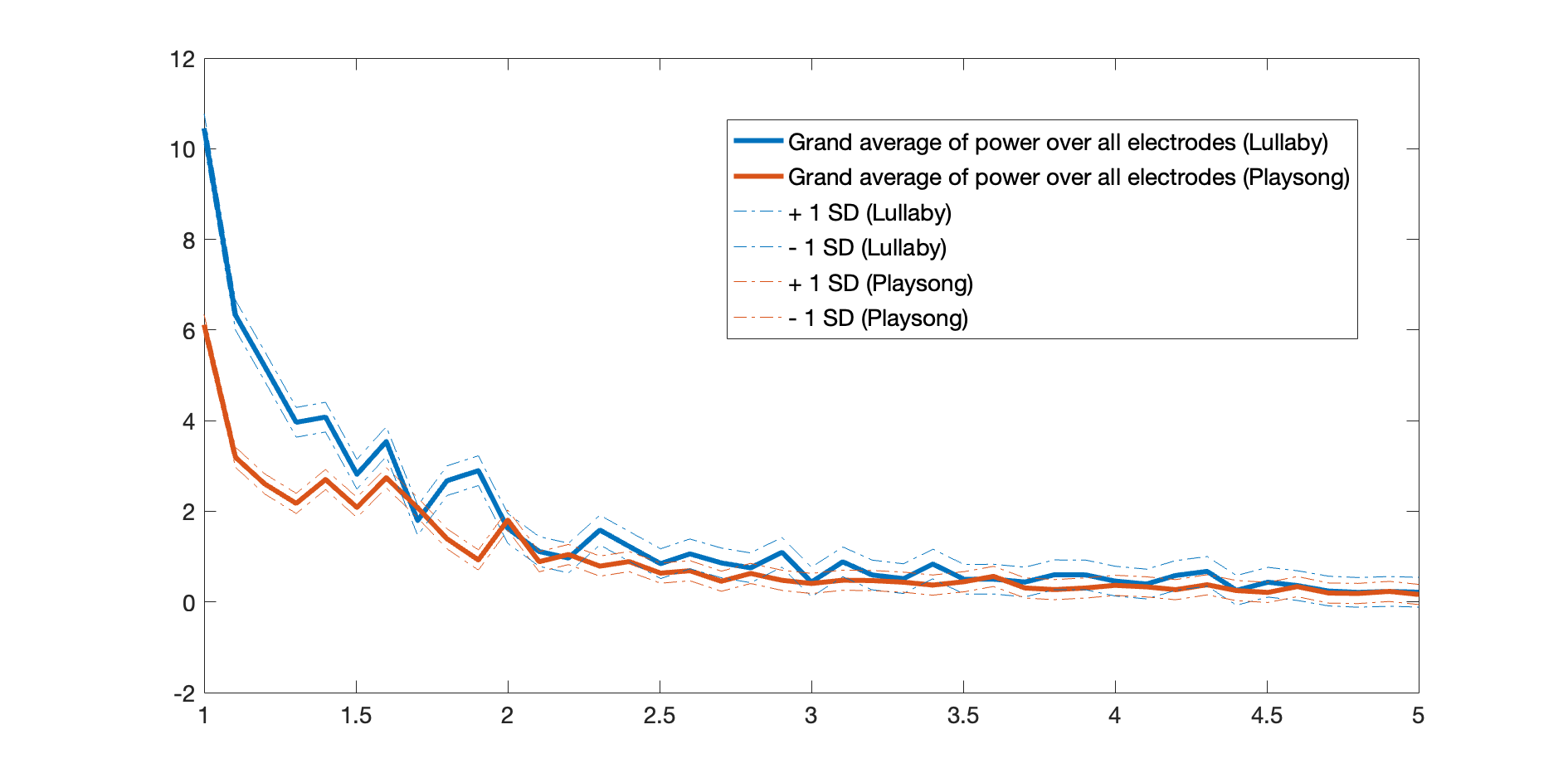

**Figure S3**. Frequency plot of infants’ EEG power averaged over each condition (lullaby = blue, playsong = red) and all channels. Frequency is plotted on the x Axis and logged absolute power is on the y Axis. EEG power in the lullaby conditions shows a slight peak over the beat-related frequencies (averaged at 2.3 Hz), individual visualization of infants' EEG power, however, shows vast individual differences mirroring the individual differences in the acoustic features of the ID songs.

**Temporal response functions from “forward” models**

We computed the individual temporal response functions (TRF) for the maternal singing envelope in the *playsong* and *lullaby* conditions in *forward* models (see *Figure S3*). TRF were compared against zero. However, we found no significant clusters in the TRF. We assume that either longer and cleaner recordings or more infants are needed to extract a clear temporal response function (Crosse et al., 2021).

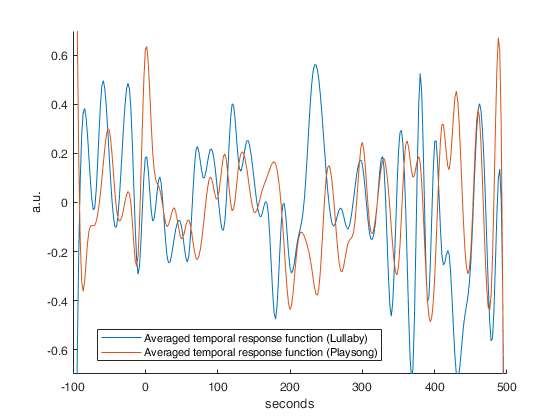

**Figure S4**. Temporal response functions of infants’ EEG signal in frontal channels in the lullaby (blue) and playsong (red) condition. Time is depicted on the x Axis, and the weights are depicted on the y Axis. The weights were neither significantly different from zero nor between each other, *p* > .30.
